# A neural decision signal during internal sampling from working memory

**DOI:** 10.1101/2022.03.31.486618

**Authors:** Freek van Ede, Anna C. Nobre

## Abstract

How humans transform sensory information into decisions that steer purposeful behaviour is a central question in psychology and neuroscience that is traditionally investigated during the sampling of external sensations. The decision-making framework of gradual information sampling toward a decision has been proposed also to apply when sampling internal sensory evidence from memory. However, neural evidence for this proposal remains lacking. Here we report that sampling internal visual representations from working memory elicits a scalp-EEG potential associated with gradual evidence accumulation – the Central Parietal Positivity (CPP). Consistent with an evolving decision process, we show how this signal (i) builds up to, and scales with, the time participants require to reach a decision about the cued memory content and (ii) is amplified when having to select (decide) among multiple contents in working memory. These results bring the electrophysiology of decision making into the domain of working memory and suggest that variability in memory-guided behaviour may be driven (at least in part) by variations in the sampling of our inner mental contents.

## Introduction

How humans transform sensory information into decisions that steer purposeful behaviour is a central question in psychology and neuroscience (1–6). Traditionally this form of decision making is studied using sensory stimulus streams containing noisy or mounting evidence, which is thought to be gradually accumulated and integrated until a decision is reached that then guides behaviour (7–11).

Like perception, working memory also serves to link relevant (memorised) sensations to purposeful behaviour (12–18), and performance often involves deciding among multiple available representations (19, 20). Accordingly, it has been proposed – on the basis of both theoretical grounds (21) and behavioural observations (22, 23) – that the decision-making framework of gradual information sampling toward a decision may also apply when sampling internal sensory representations from memory. Yet, neural evidence for this proposal has remained lacking.

Here we fill this gap by studying the Central Parietal Positivity (CPP) – a scalp EEG potential that has been shown to track gradual decision formation in humans (10, 24–28). Previous CPP studies have required participants to make decisions based on external sensations (10, 24–28). Instead, here we report a CPP-like signal when participants reached decisions based on internal evidence in visual working memory, in the absence of any external sensory evidence. Consistent with a gradual decision process, we show how this signal: (i) builds up to, and scales with, the time participants require to initiate their working-memory-guided report and (ii) is larger when having to select (decide) among more than one representation in working memory.

## Results

Participants held two tilted coloured bars in visual working memory until a colour change of the central fixation cross cued them to select and report the precise orientation of the colour-matching bar from memory (**Fig. 1a**; see also (29) for a full task description and complementary results on this task). Analysis of EEG activity evoked by the memory cue revealed a pronounced positive potential (**Fig. 1b** left, cluster P < 0.0001) with a clear central-posterior topography (**Fig. 1c**, left). This potential is reminiscent of late positive potentials previously noted in working-memory tasks (30–32). Our task, however, had the distinguishing feature that we used a symbolic cue (fixation-cross colour change) to direct attention to internal working-memory content. This is different from traditional working-memory tasks in which decisions are made in the presence of a probe stimulus to which internal memory contents have to be compared (such as in change-detection and match-to-sample tasks). In our task, decisions about how to act did not concern the cue itself, but rather the orientation of the internal memory content that was indexed by the cue.

**Figure 1.**
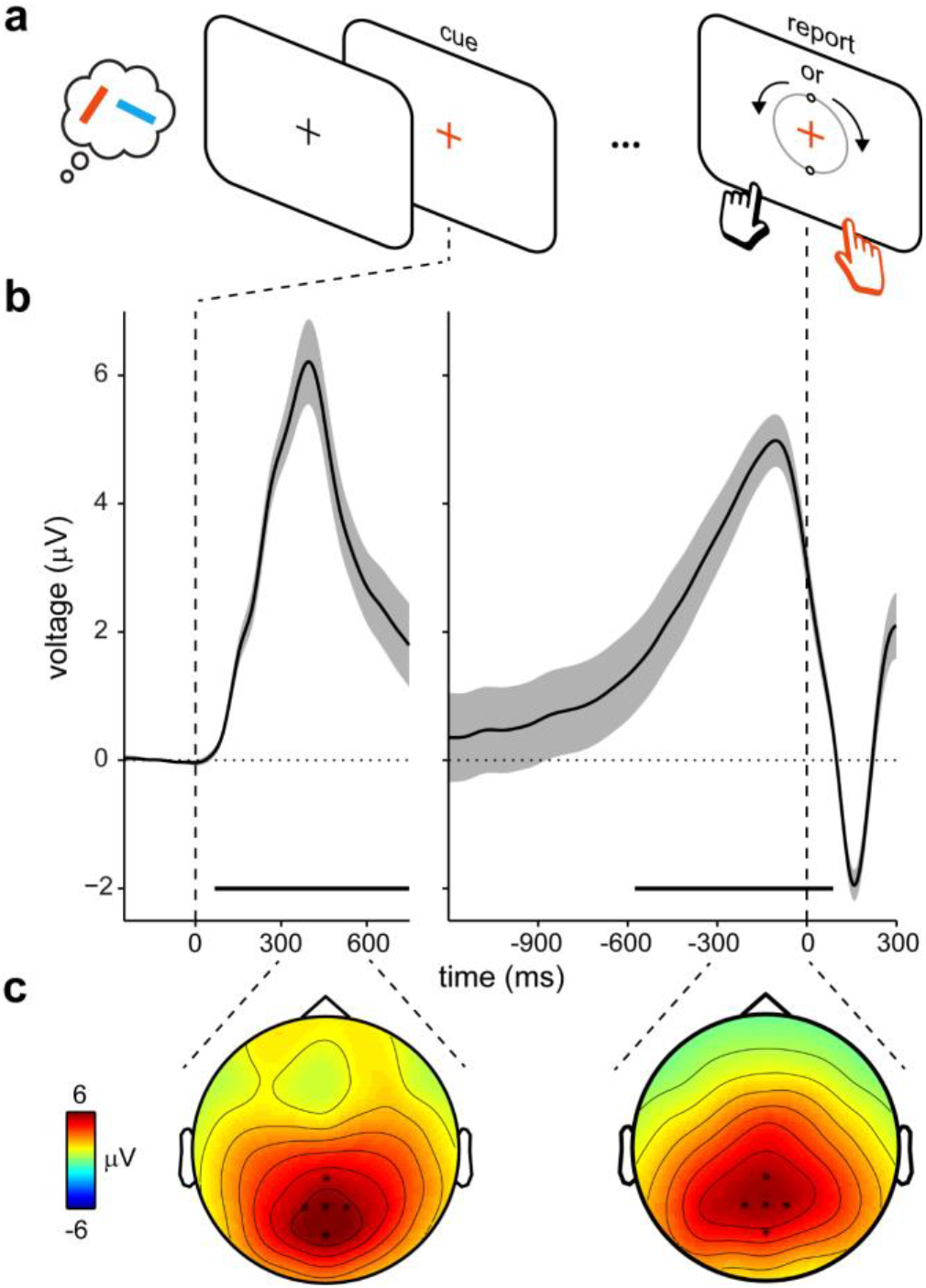
A CPP decision signal during internal sampling from visual working memory. **a)** Visual working-memory task with a cued orientation-reproduction report (adapted from (29)). Participants held two coloured oriented bars in working memory until a colour change of the central fixation-cross cued them to report the orientation of the colour-matching item from memory. The reporting dial appeared only upon report initiation (**Fig. 2a** and **Supplementary Fig. 2** for the distribution of decision times in this task). **b)** Average EEG potentials in the selected central-parietal electrodes (indicated in **c**) relative to memory-cue onset (left) and report (decision) onset (right). Shading denotes ± SE, calculated across participants (n=25). **c)** Topographies associated with the EEG potentials in **b**, averaged over the indicated time ranges (300-600 ms after cue and 300-0 ms before decision).

Aligning these data to the moment at which participants decided to initiate their report revealed a clear gradual build-up of the potential toward this decision (**Fig. 1**, right; cluster P < 0.0001). This occurred despite no additional or changing sensory inputs in this decision period (the reporting display in **Fig. 1a** appeared only at report initiation). This build-up strongly resembles an accumulation-to-bound CPP signal similar to prior studies that have attributed this component to a gradual decision signal (10, 24–28). This CPP-like response was invariant to the memorised side of the cued memory item or the hand required for reporting (**Supplementary Fig. 1**). These data thus reveal a CPP-like decision signal when sampling a cued internal representation from working memory in service of behaviour. Accordingly, and for convenience, we will refer to this signal as the CPP_i_ (“i” for internal).

To test whether the observed CPP_i_ reflected a decision-like process, we took advantage of the variability in decision times – the time from cue onset to report onset. We observed considerable variability in decision times (**Supplementary Fig. 2**), despite the relatively simple nature of our task. What may account for this variability in decision times when sampling from internal stores? We found that the CPP_i_ varied systematically with decision time over the full range of values (**Fig. 2a**). Confirming a strong correspondence between this signal and decision times, faster (slower) decisions were associated with a steeper (shallower) build-up of the CPP_i_ (**Fig. 2b**) – akin to what has been reported when deciding about external sensations (10, 24, 26–28). This relation between CPP_i_ slope and decision time (as quantified in **Fig. 2c**) was highly robust across participants (average correlation between slope and decision-time bin: r = −0.725 ± 0.082 [M ± SE]; group-level evaluation: t(24) = −8.858, p = 4.967e-09, d = −1.772).

**Figure 2.**
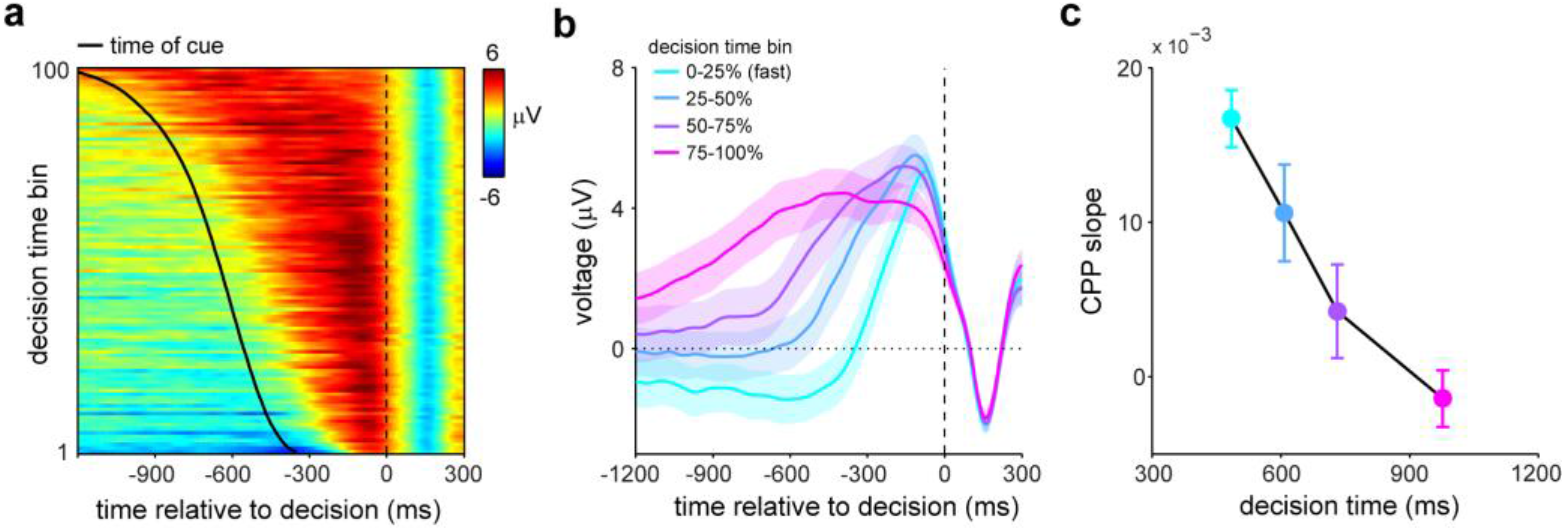
The CPP_i_ during internal sampling scales with decision time in an accumulation-to-bound fashion. **a)** Average EEG potentials in the selected central-parietal electrodes locked to report onset as a function of decision time. Data for each participant were binned into 100 percentile bins, and binned data were subsequently averaged across participants. The black line denotes average decision time in each bin. **b)** Overlay of average EEG potentials across quartile bins of decision time. **c)** Slopes associated with the data in **b**, calculated in the −500 to −50 ms pre-decision interval. Bins are positioned on the x-axis according to the average decision time in each bin. Shading in **b** and error bars in **c** denote ± SE, calculated across participants (n=25).

Finally, we asked whether this CPP_i_ signal during internal sampling depended on the requirement to select (decide) among multiple contents maintained in working memory. To this end, we re-analysed the data from a similar experiment (of which the complementary results are reported in (17)) that differed in one key aspect. In this experiment, trials contained pre-cues presented prior to memory encoding that could either be informative or neutral. In trials with a neutral pre-cue, participants still had to select the relevant memory item at the end of the memory delay (as in experiment 1; blue condition in **Fig. 3**). However, in trials with informative pre-cues, participants could pre-select the relevant memory item during initial memory encoding. Therefore, in these trials, only one item had to be retained in working memory and there was no longer a need to decide among more than one memory representation following the report cue at the end of the memory delay (red condition in **Fig. 3**).

**Figure 3.**
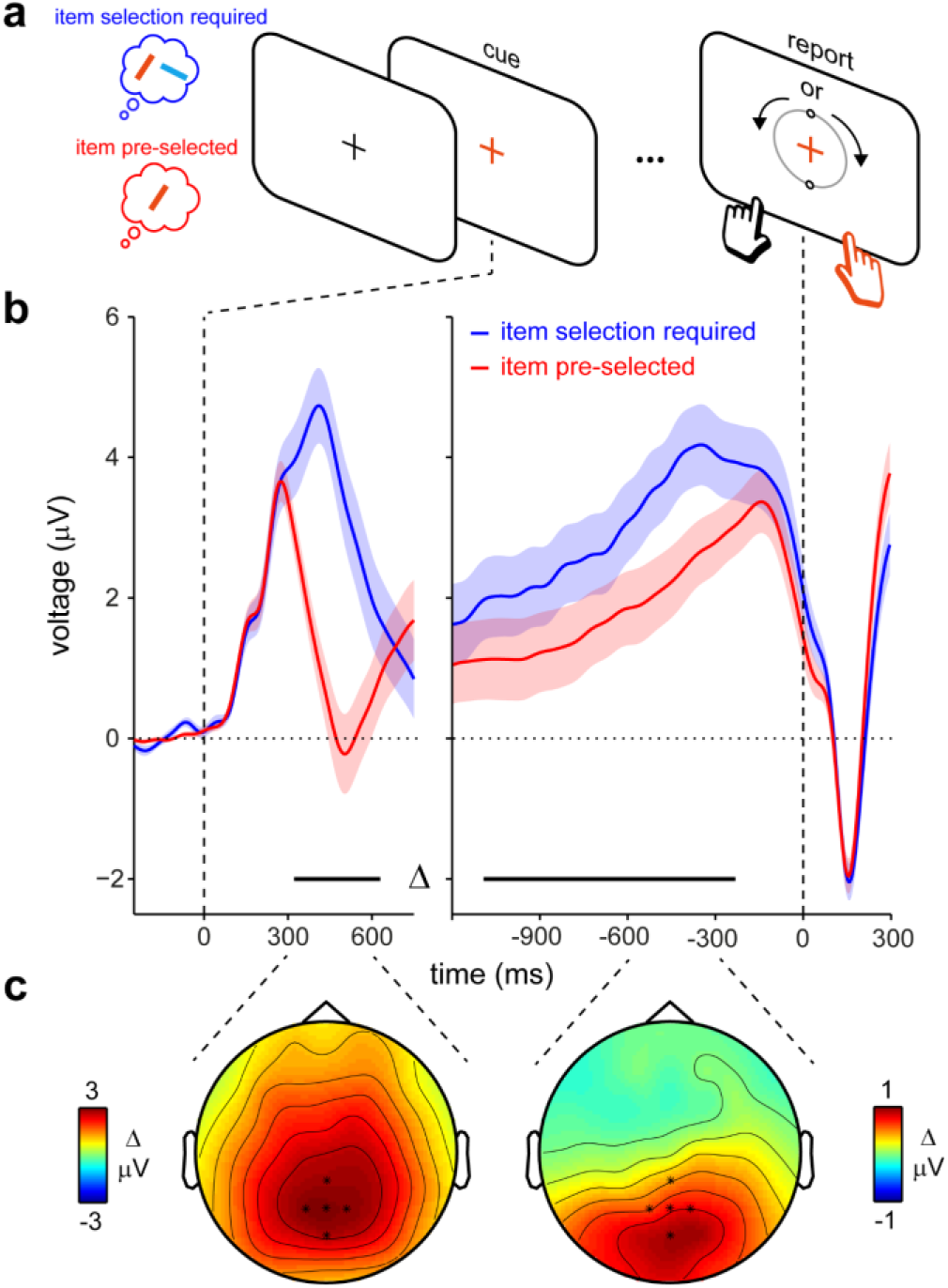
The CPP_i_ decision signal is larger when having to select among alternative working-memory contents. **a)** Visual working-memory task with one (no item-selection required) or two (item-selection required) items held in working memory prior to the memory cue (adapted from (17)). The initial encoding displays (not shown) always contained two items, of which one could be pre-cued, reducing the effective memory load to 1 (red condition). **b)** Average EEG potentials in the two conditions in the selected central-parietal electrodes relative to cue onset (left) and report (decision) onset (right). Horizontal lines denote significant differences between conditions (cluster-based permutation). Shading denotes ± SE, calculated across participants (n=25). **c)** EEG topographies associated with the condition differences in **b**, averaged over the indicated time ranges (300-600 ms after cue and 600-300 ms before decision).

In this separate dataset, we replicated our main result of a CPP_i_ during internal sampling for working memory (**Fig. 3b**). Furthermore, these data revealed that this EEG potential was significantly larger when participants were required to select among competing memory representations in the decision period between cue and report initiation (blue vs. red lines in **Fig. 3b**; topographies associated with these differences in **Fig. 3c**). This occurred both when aligning the EEG data to the report cue (**Fig. 3b**, left; cluster P < 0.0001) and to the decision (**Fig. 3b**, right; cluster P = 0.0026). Note that the difference between conditions (in the magnitude of the CPP_i_ component) was observed despite the fact that the memory report cue as well as the ensuing reporting display and reporting requirements were identical. This difference between conditions that differed solely in effective memory load must therefore be attributed to the internal cognitive processes – the selection and sampling of the relevant internal representation – in the interval between the memory cue and the initiation of appropriate memory-driven behaviour.

## Discussion

It has been posited that the decision-making framework of gradual evidence accumulation toward a decision may apply not only when sampling external sensations, but also when sampling (“retrieving”) internal sensory representations in (working) memory (21–23). Yet, neural evidence for this proposal has remained lacking. Here we fill this gap. Building on the established link between the CPP and gradual decision formation during perceptually guided behaviour (10, 24–28), our CPP_i_ results suggest that similar neural decision processes may support memory-guided behaviour.

Our results link variability in the speed of memory-guided behaviour to variations in the sampling of internal memory contents – as reflected in the CPP_i_. This type of variability is not typically the focus in studies on visual working memory, which tend to focus on mechanisms of memory retention (during the delay) instead of utilisation (after the delay). Yet, this type of variability is highly relevant from the perspective that working memory – just like perception – ultimately serves to translate (past) sensations into purposeful (future) behaviour (12, 14–19). If we consider that internal memory representations are inevitably subject to noise (33–36) and may vary in their precision (37), an internal evidence accumulation process may ensure optimal translation of noisy internal sensory representations into adaptive behaviour.

The notion of gradual sampling of relevant sensory information toward a decision to guide behaviour has a strong tradition in psychology and neuroscience (1–11). To date, however, this form of decision making has been studied almost exclusively for decisions that regard currently available sensory information – even if it has been appreciated that memory processes often play a key role in reaching the best decision in such situations (38–40). Our data suggest that similar gradual decision formation may also apply in cases where the decision itself regards the visual identity of an *internal* memory representation. This brings together the study of working memory and decision-making – two topics that are often considered separately, despite their common focus on how the brain translates sensory information into purposeful behaviour.

Out data also build on studies of working-memory retrieval that have previously implicated a role for the P3 EEG potential (30–32), which is related to the CPP (27). Specifically, we reveal the ramping nature of this EEG potential (when aligned to report onset) and demonstrate this in the absence of any external stimulation contributing evidence to weigh into the decision. This is different from traditional working-memory tasks in which decisions are made in the presence of a probe stimulus to which internal memory contents have to be compared (as in change-detection and match-to-sample tasks). This unique aspect of our task is also relevant when relating our CPP_i_ findings to prior studies reporting a gradual (decision-locked) CPP in tasks with isolated perceptual events (27, 28). In contrast to these studies, in which the decision regarded the perceptual stimulus itself, the decision in our task regarded memorised item orientation; a sensory feature exclusively contained in memory.

We found that the CPP_i_ was larger when participants had to select among two visual items in working memory, compared to when the relevant memory item had already been pre-selected (contrary to prior reports of *smaller* probe-locked P3 responses with higher memory loads (30, 31)). There are at least two possible interpretations for this CPP_i_ amplification (which are not mutually exclusive). First, the process of selecting the correct memory item could itself be conceived of as a decision (that complements the decision about the required action given its orientation). Indeed, deciding among competing alternatives (in addition to the appropriate course of action) is a central component of decision making (38, 41–43). Second, when the memory item that will become relevant is known in advance, planning for the appropriate course of action can start prior to the memory cue (17, 44). This may alleviate elements of the decision-making process that would otherwise be possible only after the memory cue, such as deciding what hand to use for responding.

We have cast our results from a gradual decision-making perspective. Could our results also be framed in terms of action preparation, without additionally invoking the notion of a “decision process”? In our task (as in most perceptual decision-making tasks), decisions were inherently linked to actions as they directly informed the to-be-initiated manual report. Yet, a pure action-preparation account with no decision component may be insufficient to capture relevant aspects of our data. First, we found a gradual build-up of the CPP_i_, whose slope varied systematically with decision time. Thus, variability in the onset of memory-guided behaviour was linked directly to variability in the gradual build up toward the appropriate action (a key indicator of a decision process). For comparison, imagine an alternative scenario in which variability in the onset of memory-guided behaviour would be due exclusively to variability in occasional ‘lapses of attention’ after the memory cue. Such a scenario would yield a decision-locked EEG signal that would be invariant to the time leading up to the report. Second, we found a larger CPP_i_ when participants were required to decide among more than one content in memory, despite equivalent reporting demands in both cases. Finally, our CPP_i_ had a characteristic central-posterior topography (linked to the parietal cortex (30) by a prior EEG-fMRI study), rather than a more frontal distribution that would be expected from a source in (pre)motor areas. This topography was furthermore invariant to the hand required for reporting, in agreement with other studies that have shown that the CPP can be dissociated from effector-specific action planning (25, 45).

Taken together, our results bring the electrophysiology of decision making into the realm of working memory and suggest that variability in memory-guided behaviour may be accounted for (at least in part) by variations in the sampling of our inner mental contents.

## Methods

The current article presents the re-analyses of data from two prior studies (“E1” and “E2” for convenience), which each used a similar task set-up. Complementary results from both prior studies were published previously (E1: (29); E2: experiment 1 in (17)). These prior publications focused on distinct questions regarding the relation between visual working memory and action planning. Instead, we here report analyses prompted by contemporary studies on the Central Parietal Positivity (CPP) as a human EEG signature of perceptual decision-making (10, 24–28). Our task designs and data proved ideally suited to address whether a CPP-like signature could also be established when human participants decide how to act based on internal representations in working memory.

### Participants

E1 contained data from 25 healthy human volunteers (14 female; age range 19-36; 2 left handed), as did E2 (18 female; age range 18-35; all right handed). Sample sizes of 25 were set a-priori as described in (17, 29). Participants provided written informed consent before participation and received £15/hour for their participation. Experimental procedures were reviewed and approved by the Central University Research Ethics Committee of the University of Oxford.

### Task design and procedure

In both E1 and E2, participants performed a visual working memory task with a delayed orientation-reproduction report from memory.

In E1 (see also (29)), participants encoded two coloured tilted bars (presented for 250 ms) into working memory. After a delay (drawn randomly between 2000-2500 ms) in which only the fixation cross remained on the screen, participants were cued to select one of the two memory items for report. The cue consisted of a colour change of the central fixation cross. The task was to reproduce the orientation of the memory item that matched the colour of the cue. At encoding, the two bars were positioned approximately 5.7 degrees visual angle to the left and right of fixation. Bars could be either of four clearly distinguishable colours (blue, orange, green, purple) and could range in orientation between ±20 to ±70 degrees. Bar location, colour, and orientation varied independently across trials.

The orientation-reproduction report was performed by rotating a reporting dial (presented centrally, around fixation) to the memorised orientation of the cued item. The report was initiated by pressing either of two keys on the keyboard (‘\’ or ‘/’) that were operated with the left and right index finger respectively. The dial appeared on the screen once either key was pressed. Upon key press, the dial would start in vertical position and would rotate leftward (key ‘\’) or rightward (key ‘/’) until key release, which terminated the response. The dial could be rotated maximally 90 degrees. As a consequence, the tilt of the bar was directly linked to the hand required for responding: a left-tilted bar required a left-hand response, while a right-tilted bar required a right-hand response (see (29) for further details behind the rationale of this manipulation).

E2 (see also (17)) relied on the same basic task as E1 with two additions, of which the second is relevant to the current investigation. First, in E2, the memory delay was either two or four seconds (as motivated in (17)). This aspect was not of interest here, and we therefore collapsed trials with either delay duration. Second, and of direct interest, in E2 we included trials with a brief (250 ms) colour pre-cue that occurred one second *before* the encoding of the two memory items. Pre-cues informed which item would be tested later. Accordingly, in these trials (comprising 80% of all trials in E2), participants needed to retain only a single item during the working-memory delay. Consequently, they no longer needed to select the relevant memory item before initiating their memory-guided report (which was still prompted by the second colour cue at the end of the memory delay). In the remaining 20% of the trials the pre-cue was neutral (grey) and participants were required to keep both items in working memory until the coloured report cue indicated the relevant item to reproduce at the end of the delay (as in E1). This enabled us to compare the EEG response in the period between the reporting cue and the decision in trials in which the relevant memory item (a) had already been selected prior to the reporting cue or (b) still required to be selected after the report cue. Importantly, the memory report cue itself, as well as the ensuing reporting display and response requirements were identical between these two conditions.

In both E1 and E2, participants completed two consecutive EEG sessions of approximately 50 minutes. Sessions were broken down into ten blocks of approximately five minutes. In E1, each block contained 60 trials, whereas in E2 each block contained 40 trials. This yielded 1200 trials per participant in E1 and 800 in E2.

A critical feature of both E1 and E2 was that we probed (cued) internal working-memory content using a mere colour change of the central fixation cross, without presenting any new visual element on the screen until response initiation (at which point the reporting dial appeared). For the current purposes, this is an important advantage over common working-memory tasks in which a perceptual ‘probe stimulus’ is presented after the delay, to which a memory representation must be compared (such as in change-detection or delayed match-to-sample tasks). In such cases, the probe stimulus may itself trigger perceptual decision-making and yield a perceptually driven CPP. In contrast, in our tasks the only perceptual element that cued the decision-making process was a colour change of the central fixation cross. Crucially, while the colour change was a perceptual element, our task was not to decide about colour, but rather to sample the appropriate orientation information from working memory for the orientation-reproduction report. Colour served merely as a medium to cue the correct internal representation. Moreover, because possible cue colours were highly distinguishable (blue, orange, green, purple) there were minimal demands on any “decision process” regarding perceived colour itself.

Another key feature of our task was that once a key press was initiated there was no return. The report terminated at key release, without opportunity for further adjustment. Because of this, participants were required to deliberate carefully before committing to the initiation of their report.

### EEG acquisition and preprocessing

EEG acquisition and analyses followed the same procedures in E1 and E2. EEG data we collected using SynAmps amplifiers and Neuroscan software (Compumedics). Electrode placement followed the international 10-10 system, using a 61-electrode montage. Data were referenced to the left mastoid at acquisition. Measurements of vertical and horizontal EOG were included for offline ICA correction. Data were digitised at 1000 Hz.

EEG data were analysed in Matlab, using the FieldTrip toolbox (46). Data were re-referenced to the average of both mastoids, and down-sampled to 250 Hz. ICA was performed to identify components associated with blinking or lateral eye movements and bad components were removed from the data. After ICA correction, trials with exceptional variance were identified based on visual inspection (ft_rejectvisual with the ‘summary’ method) and excluded. Noisy trials were identified without knowledge of the relevant conditions to which particular trials belonged. We additionally excluded trials in which the decision time (the time between cue onset and report onset) was below 200 or above 2000 ms. In E1, on average 1027 ± 25 trials (86 ± 2 %) were retained for analysis. In E2, on average 700 ± 11 trials (87 ± 1 %) were retained for analysis.

### CPP analyses

To target the CPP, we zoomed in on an a-priori defined electrode cluster centred on electrode Pz (‘Pz’, ‘CPz’, ‘POz’, ‘P1’, ‘P2’), consistent with prior work investigating the P3 and CPP in the context of perceptual decision-making (e.g. (27)).

To characterise the CPP, we aligned all trials to two moments: (1) the onset of the colour cue at the end of the memory delay, which starts the decision-making process, and (2) the onset of the memory-guided reproduction report or “decision”, which signals the completion of the decision-making process. We focused our primary analyses on the decision-aligned data because the EEG signal in this alignment is less dominated by the sensory-driven ERP evoked by the abrupt colour change of the cue. As such, the decision-aligned data provides a cleaner evaluation of the gradual build-up response to the decision – a key defining feature of the CPP (e.g. (27)).

We baseline corrected the cue-locked data by subtracting the average EEG signal in the 250 ms window prior to cue onset. Likewise, we baseline corrected the decision-locked data by subtracting the average EEG signal in the 250 ms window following response initiation. Baselining ensured that the trials were comparable with regard to the event to which we aligned the data, regardless of trial-to-trial variations in the time between cue onset and decision onset. As an alternative to low-pass filtering, CPP time courses were smoothed with a Gaussian kernel (with a standard deviation of 30 ms).

To evaluate the relation between the CPP and decision times (defined as the time between cue onset and report initiation) we first sorted the data of each participant into non-overlapping bins of decision times and calculated the average ERP in each bin. To leverage the large number of trials we had available in E1, we performed this analysis using 100 percentile bins per participant. After binning, we averaged the binned data across participants, and visualised these as an ERP image (47), sorted by decision-time (as in (10)). To facilitate additional visualisation and quantification, we also performed the same analysis using four quartile decision-time bins per participant.

### Statistical evaluation

We used a cluster-based permutation approach (48) to evaluate the baseline-corrected trial-averaged EEG potential time courses against zero (E1) and well between experimental conditions (E2). The cluster-permutation approach enabled us to evaluate statistically our data along the relevant time axes, while circumventing the multiple-comparison problem.

To evaluate the link between the identified CPP and decision times, we compared the slope of the CPP across the four quartile bins of decision times. For the data from each quartile, we quantified CPP slope as the linear regression coefficient between time and the trial-average EEG potential in the [−500 to −50] ms window relative to response initiation. We quantified the gradient in this slope across the four quartile bins using a Pearson’s correlation between bin number and slope coefficient. We obtained this correlation for each participant (first level) and quantified this at the group level (second level) using a one-sample t-test against zero (with zero denoting the null-hypothesis of no change in CPP slope across the four decision-time bins).

We restricted our statistical analyses to the EEG time courses extracted from the a-priori defined electrode cluster. We additionally included topographical analyses to visualise the reported effects. Topographies were not subjected to additional inferential statistical testing. Instead, their visualisation served primarily to increase transparency, to reinforce the suitability of our a-priori electrode selection, and to demonstrate the ‘plausibility’ of the reported findings (49).

### Data availability

Data from both experiments have been made publically available previously. Data from E1 are available through the Dryad Digital Repository (at https://doi.org/10.5061/dryad.sk8rb66). Data from E2 are available through the Zenodo Repository (at https://doi.org/10.5281/zenodo.4471943). Data from E2 in the current article are the data from “experiment 1” in the latter data package that itself contains data from two experiments.

## Acknowledgements

This research was supported by an ERC Starting Grant from the European Research Council (MEMTICIPATION, 850636) to F.v.E., a Wellcome Trust Senior Investigator Award (104571/Z/14/Z) and a James S. McDonnell Foundation Understanding Human Cognition Collaborative Award (220020448) to A.C.N, and by the NIHR Oxford Health Biomedical Research Centre. The Wellcome Centre for Integrative Neuroimaging is supported by core funding from the Wellcome Trust (203139/Z/16/Z). The authors also wish to thank Sammi Chekroud, Sage Boettcher, and Daniela Gresch for their contributions to the research in the original publications, and Méadhbh Brosnan and Rose Nasrawi for valuable discussions during the preparation of the manuscript.

This research was funded in part by the Wellcome Trust [Grant numbers 104571/Z/14/Z, 203139/Z/16/Z]. For the purpose of open access, the author has applied a CC BY public copyright licence to any Author Accepted Manuscript version arising from this submission.

## Supplementary Material

**Supplementary Figure 1.**
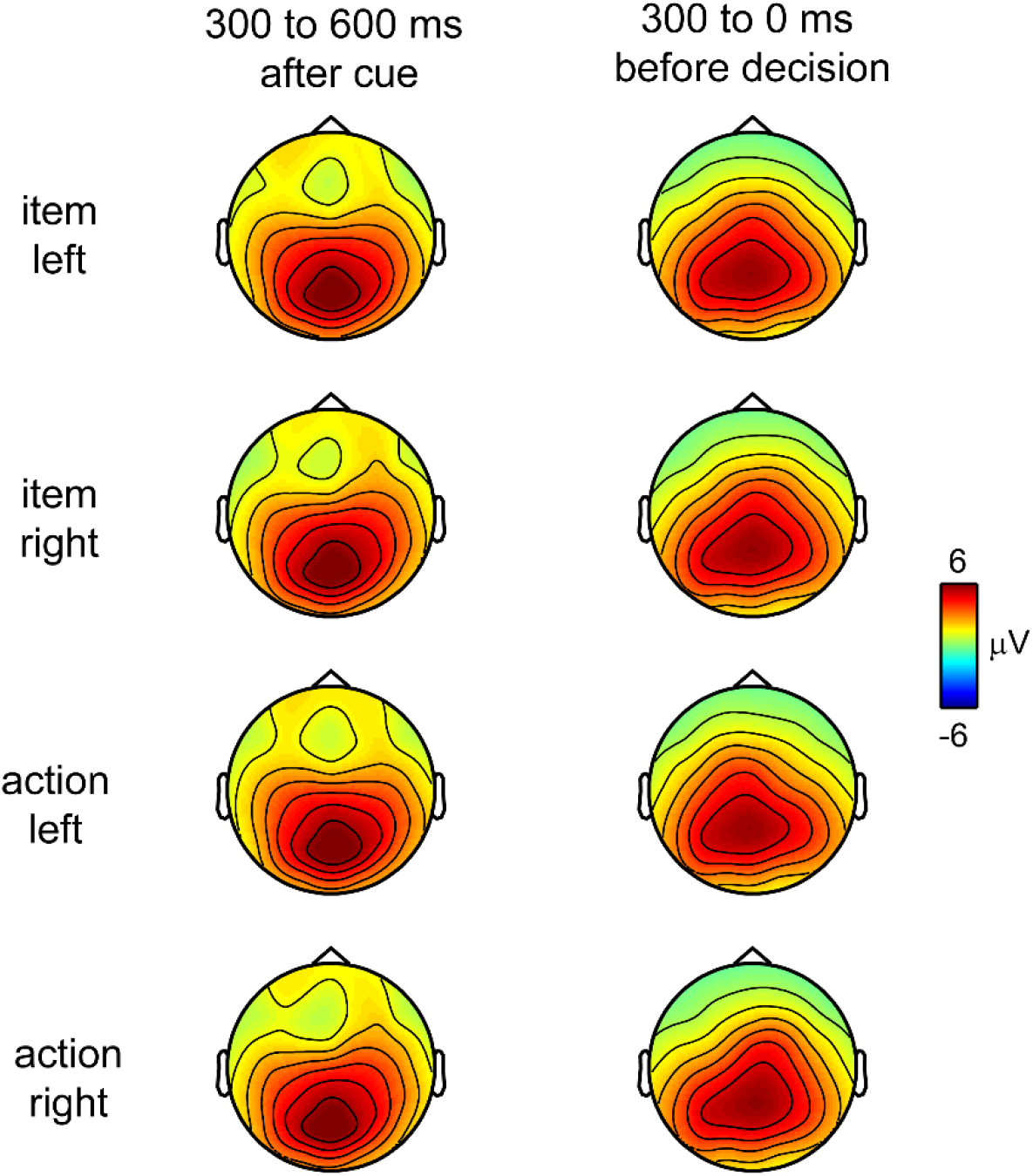
The CPP_i_ topography is invariant to the visual location and the manual action associated with the cued memory item. CPP_i_ topographies relative to cue (left) and decision (right) as a function of whether the cued memory content (item) was the left or right item at encoding, and whether the reproduction report (action) required the left or the right hand. Data from Experiment 1.

**Supplementary Figure 2.**
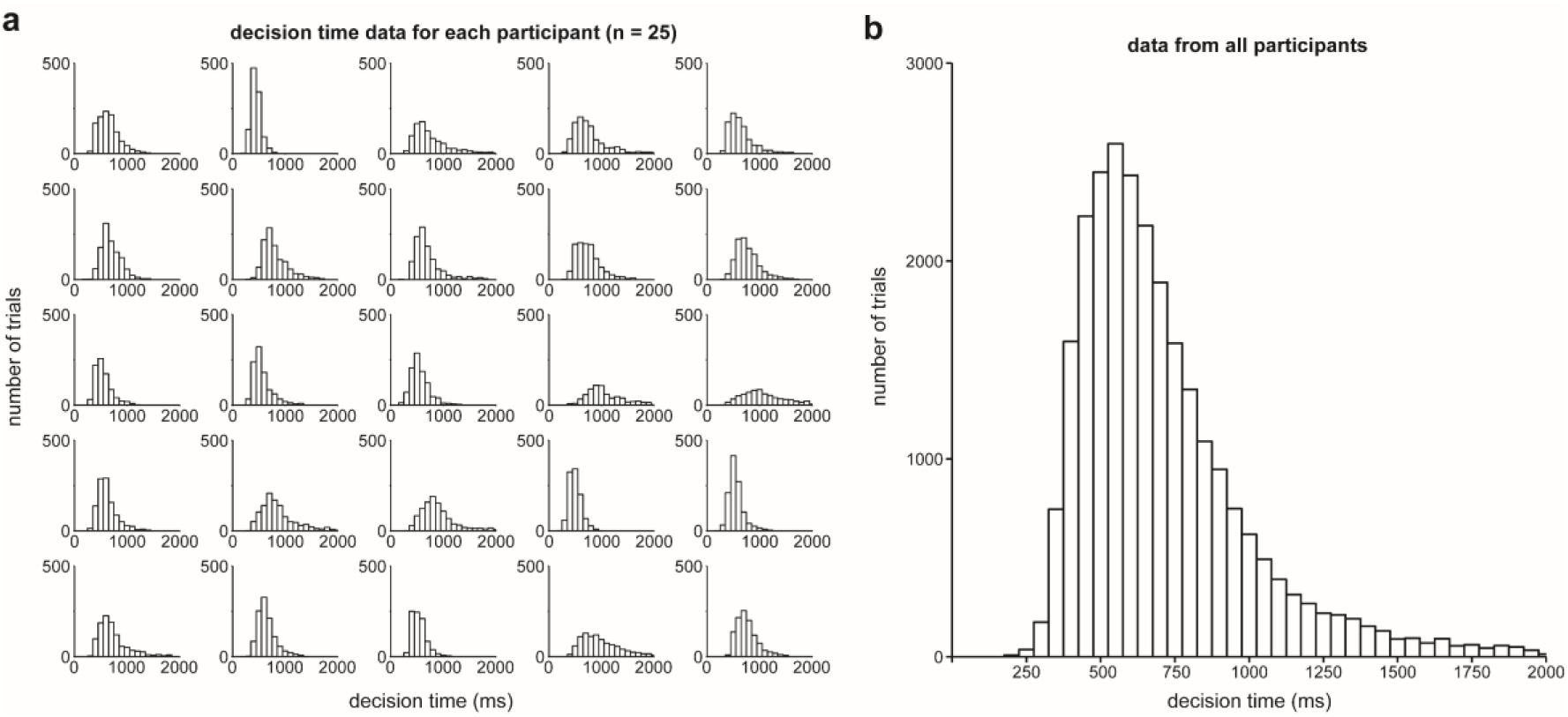
Decision-time distributions. **a)** Decision time distributions across trials for all individual participants in experiment 1 (n=25). **b)** Group-level decision-time distribution, aggregating decision times across trials and participants. Trials in which the decision time exceeded 2000 ms were excluded from further analysis.

